# Micromorphology and ammonium transmembrane transport characteristics in roots of rice seedlings

**DOI:** 10.1101/2025.01.07.631647

**Authors:** Xiaoguo Zhang, Yanan Xu, Song Chen, Guang Chu, Chunmei Xu, Yongjie Yang, Danying Wang

## Abstract

During the seedling stage, rice root exhibit rapid growth and absorb significant quantities of nutrients. This study involved cultivating rice seedlings in two different seedling-raising mediums characterized by high ammonium concentration (HN, NH_4_^+^ 1.2 mg g^−1^) and low ammonium concentration (LN, NH_4_^+^ 0.006 mg g^−1^). The morphology of distinct root zones (root cap, meristematic zone, elongation zone, and maturation zone) was observed under a microscope throughout root development, and variations in transmembrane flux rates of NH_4_^+^ in different root zones were assessed using Non-invasive micro-test technology (NMT). Results showed that the root caps of both seminal and adventitious roots became separated from the root tip between 9 and 12 days after root emergence. During this time, the root tip transformed from a sharp to a round shape, and the distance from the maturation zone to the root tip progressively decreased from over 900 μm on 6 days after sowing (DAS) to under 500 μm on DAS12. The meristematic zone was the primary site for NH_4_^+^ absorption in new root and proved to be particularly sensitive to environmental NH_4_^+^ concentration. At DAS 6, the net NH_4_^+^ flux rate was highest in the meristematic zone under both HN and LN treatments. Additionally, the external NH_4_^+^ concentration influenced the direction of NH_4_^+^ flux in the meristematic zone at DAS9, with HN being a net NH_4_^+^ influx and LN being an NH_4_^+^ efflux, and the net NH_4_^+^ influx in the meristematic zone persisted for 3 days longer in the HN treatment compared to the LN treatment. In mature roots, the root hair zone emerged as the primary site of NH_4_^+^ uptake, exhibiting an infNH_4_^+^ influx rate of 40−140 pmol cm^−1^ s^−1^, and this low rate of uptake could be sustained for up to 12 days after root emergence. By DAS15, a net NH_4_^+^ efflux was observed in the entire seminal root, signaling the loss of NH_4_^+^ absorption function 15 days post-emergence. A similar trend was noted in adventitious roots, where NH_4_^+^ uptake remained functional for 12-15 days after root emergence. As rice seedlings continued to grow, new adventitious roots replaced old ones, facilitating ongoing NH_4_^+^ uptake.

## Introduction

Studying the changes in root morphology and nutrient uptake capacity during the growth of rice seedlings is crucial for evaluating and cultivating robust seedlings, as the root growth and nutrient uptake capacity of rice seedlings are closely linked to their robustness. The rice root system comprises seminal root, adventitious roots and lateral roots (Coudert *et al*., 2010), with the seminal root emerging directly from the radicle during seed germination and serving as the primary supporting and absorbing tissue of rice until the 3-leaf stage (Yang and Tu, 2011; Wang *et al*., 2011). As the nutrient and water requirements of rice growth increase, adventitious roots gradually developed from the bottom tiller nodes through polar transport of growth hormones, cytokinin and other actions (Giri *et al*., 2018; Liu *et al*., 2009).

The rice root’s longitudinal anatomy can be categorized into four zones: the cap, meristematic zone, elongation zone, and maturation zone, progressing from the root tip upwards. Positioned at the top of the rice root, the root cap consists of large, irregularly arranged cells that form a protective structure surrounding the outer part of the meristematic zone; it serves as the dual purpose of safeguarding the root system and sensing external environment to relay signals (Chen *et al*., 2006; Wang *et al*., 2014). The meristematic zone is dense packed with cells due to continuous proliferation (Hayashi *et al*., 2013), and cell differentiation timing is regulated by cytokinin levels, influencing the size of the meristematic zone (Dello Ioio *et al*., 2007; Moubayidin *et al*., 2009). Above the meristematic zone lies the elongation zone, where cells begin to elongate (Wang, 2020), and further up is the maturation zone, where cells stop elongating and some of them protrude outwards to form root hairs (Du *et al*., 2018; Kong, 2009).

Functions of rice roots include fixation and support, absorption and transportation, synthesis and secretion. Among these functions, absorption is crucial for providing water and nutrients essential for rice normal growth (Meng *et al*., 2019). Previous researches on rice roots have primarily focused more on the entire plant root system, investing the correlation between morphological factors (such as root number, length and volume) and physiological indicators (including root vigor and root respiration intensity) with nutrient uptake, yield and quality formation (Chen *et al*., 2017; Liu *et al*., 2002; Ju *et al*., 2015). In certain instances, redundant root growth can occur, making it unreliable to evaluate N use efficiency solely based on root morphology and distribution (Liu *et al*., 2002). Delving into the nutrient absorption of individual root and examining the distinct functions of various root components (such as the root cap, meristematic zone, elongation zone and maturation zone) at a microscopic level could offer deeper insights into the mechanism of efficient nutrient uptake in rice.

Nitrogen (N) is a crucial element for plant growth, and NH_4_^+^ serves as the primary inorganic form of N absorbed by rice. Research indicated that rice variety exhibiting efficient N utilization possess roots had high porosity and radial oxygen secretion capacity, leading to an increase in ammonia-oxidizing bacteria and mineral N content in the rhizosphere soil under low N conditions (Chen *et al*., 2020). However, in environments with high NH_4_^+^ conditions, rice roots have a tendency to absorb excess NH_4_^+^, and a large amount of NH_4_^+^ accumulates in the roots without being fully transformed and utilized (Hao *et al*., 2020). Some of the excess NH_4_^+^ is released from the root, so there are both transmembrane NH_4_^+^ influx and efflux in rice roots (Sun *et al*., 2018; Taylor and Bloom, 1998; Verbelen *et al*., 2006; Zhou *et al*., 2003). Chen *et al*. (2013) discovered that the meristematic zone was the primary site for NH_4_^+^ transmembrane influx, while transmembrane efflux predominantly occurred in the root elongation zone. In our previous research, the rate of NH_4_^+^ transmembrane influx in the root meristematic zone in N-efficient varieties was significantly higher compared to N-inefficient varieties. Additionally, there was a notable distinction in NH_4_^+^ transmembrane flux between new and old roots within the same root zone (Zhang *et al*., 2015a). Therefore, there is an immediate necessity to investigate the variations in NH_4_^+^ uptake capacity of roots over time and the NH_4_^+^ uptake substitution between new and old roots to comprehensively analyze the mechanism of efficient N uptake in rice.

## Materials and methods

### Plant material and experimental treatments

The experiment was conducted in Fuyang, Hangzhou, Zhejiang (30°05′N, 119°90′E, 21 m altitude), in the years 2021 and 2022. Seeds of hybrid rice variety Yongyou1540 were first soaked at 30°C for 48 hours (h), followed by germination at 35°C for 24 h until the radicle emerged from the seed coat. Subsequently, they were sown on two different seedling raising mediums with high NH_4_^+^ concentration (HN, NH_4_^+^ 1.2 mg g^−1^) and low NH_4_^+^ concentration (LN, NH_4_^+^ 0.006 mg g^−1^). The HN and LN seedling raising mediums were placed in separate seedling trays, with three trays per treatment as three replications. After planting the germinated seeds, the seedling trays were placed in a glass greenhouse to receive solar radiation and maintain a temperature of 25−28□, following conventional seedling raising practices for water management. At DAS 6, DAS 9, DAS 12 and DAS 15 d, ten rice seedlings were sampled in each replication, and the roots were thoroughly rinsed with water for morphological and physiological analysis.

### Measurement and analysis

The rice seedlings were divided into root and shoot sections, and the number of roots were counted and root lengths were measured. The various root zones -including the root cap, meristematic, elongation, maturation zones and the root hair region of both seminal and adventitious roots - were observed under an inverted microscope (XY003-Y012; Xuyue (Beijing) Sci. & Tech. Co., Ltd., Beijing, China.) at magnification of 20x.

The NH_4_^+^ concentration in roots was measured by taking a precise 0.2 g sample of fresh root. Free NH_4_^+^ was extracted by grinding the sample with 10% glacial acetic acid, while 30% trichloroacetic acid (TCA) was added to inhibit nitrate reductase activity. The NH_4_^+^ concentration was determined using the phenol-hypochlorite method, with ammonium sulfate as the standard solution. The root NH_4_^+^ accumulation per plant was calculated by multiplying the NH_4_^+^ concentration by the root weight per plant.

Following the method described by Zhang *et al*. (2015b), NH_4_^+^ fluxes of the root cap, meristematic, elongation, maturation zones were assessed using Non-invasive Micro-test Technology (NMT, Younger USA Science and Technology Corp, USA). The measurements were carried out at room temperature (24−26°C). Ten representative intact roots were selected for testing in each replication. The roots were first immersed in an equilibrium solution (0.1 mM NH_4_Cl, 0.1 mM CaCl_2_, pH 6.0) for 10 min. After equilibration, the roots were transferred to a measuring chamber, a 3 cm diameter plastic dish containing 2 ml of fresh measuring solution (0.1 mM NH_4_Cl, 0.1 mM CaCl_2_, pH 6.0), and fixed in place for measurement. Two microelectrodes were positioned 5 µm and 35 µm from the root surface within the measuring solution. Background reading was taken by positioning the electrodes in the solution without roots. Glass microelectrodes with apertures ranging from 2 to 4μm were fabricated by Xuyue Science and Technology Co., Ltd. The net NH_4_^+^ fluxes were measured individually in the root cap, meristematic zone, elongation zone, maturation zone and root hair region, in each measurement lasting for 10 minutes. The final flux values for each zone were calculated as the averages of measurements from ten individual root in each replication (Supplementary Fig. S1).

### Statistical analysis

Figures were processed by Microsoft Excel in 2010 version (Microsoft, Redmond, WA, USA). Two-way analysis of variance (ANOVA) was analyzed in SAS 9.4 software (SAS Institute Inc., Cary, NC, USA), and means were subjected to Tukey’s honestly significant difference (HSD) test at the 0.05 probability level.

## Results

### Morphological characteristics of rice seedling root

As shown in supplementary Fig. S2, the seminal root was white at DAS 6 and adventitious roots started to emerge at the base of the stem. By DAS12, rice seedling had 9−17 roots, increasing to 15−18 roots at DAS 15. Based on the timing of their appearance, adventitious roots were categorized as either adventitious root □ and adventitious root □: those emerging at DAS 6 were designated as adventitious root □, while those emerging at DAS12 were classified as adventitious root □. Microscopic examination of the seedling roots showed that the root cap detached from the root tip between 9 to 12 days after root emergence (Fig. 1), causing the root tip to change from sharp to round. As the seedling roots aged, the distance from the maturation zone to the root tip gradually decreased. For example, the distance from the maturation zone to the tip of the seminal root was 900−1200 µm at DAS 6, reducing to 700−1000 µm at DAS 9, and further shortened to 200−400 µm at DAS 15. A similar trend was observed for adventitious roots, with the distance from the maturation zone of adventitious root I to the tip being 1200−1500 µm at DAS 6, decreasing to approximately 70% of this length after 6 days.

**Fig. 1.**
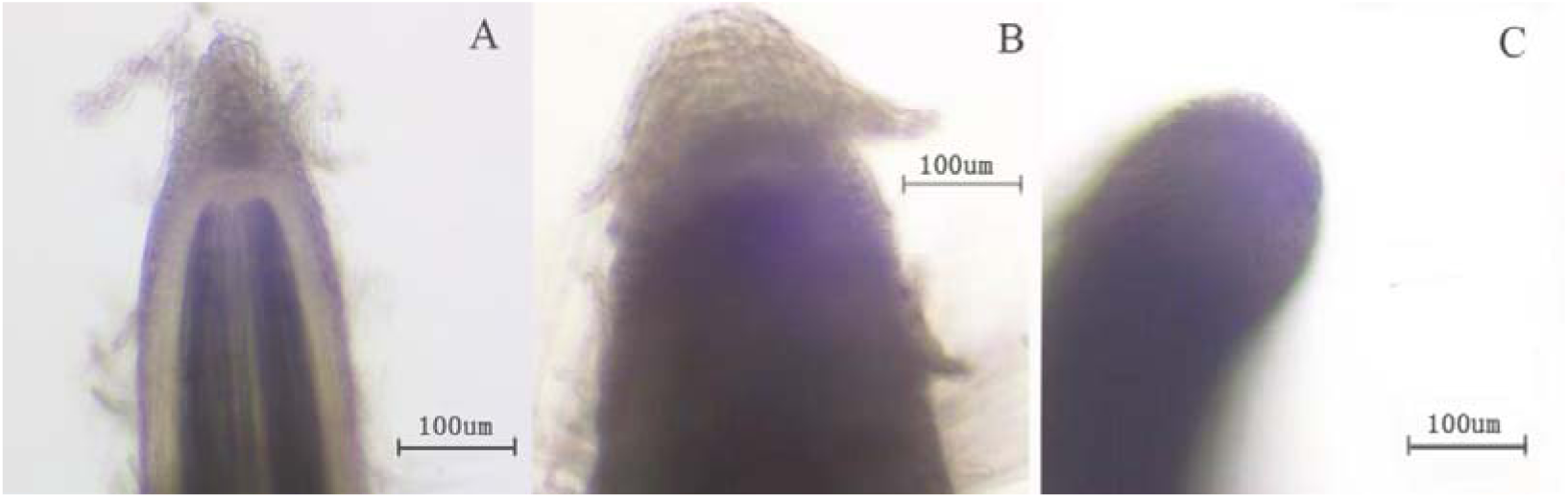
The process of root cap detachment (A before detachment at DAS6; B during root cap detachment at DAS9; C after detachment at DAS12).

### Plant NH_4_^+^ concentrations and accumulation

The root NH_4_^+^ concentrations in both HN and LN treatments showed significant variations from DAS 6 to DAS 15 (Fig. 2a). The NH_4_^+^ concentration in the seminal roots peaked at 9.9−10.3 µM g^−1^ fresh weight (FW) at DAS 6, then rapidly to decline to 3.0−3.5 µM g^−1^ FW at DAS 9, increased to 5.8−6.1 µM g^−1^ FW at DAS 12, and dropped again to 2.4−2.6 µM g^−1^ FW at DAS 15. Despite this fluctuation, the NH_4_^+^ accumulation of roots (Fig.2b) and the fresh weight of roots and shoots (Fig. 2c, 2d) continued to rise from DAS 6 to DAS 12. The varying NH_4_^+^ concentration in roots might be attributed to the mismatch between root NH_4_^+^ absorption capacity and the increase in biomass. At DAS6, adventitious roots were just starting to grow in the seedlings, and nutrients were primarily derived from the seeds. Consequently, there was no significant disparity in NH_4_^+^ concentration in roots between the HN and LN treatments. From DAS6 to DAS9, NH_4_^+^ in root was rapidly assimilated and utilized as seedling developed, leading to a decline in NH_4_^+^ concentration in the roots. However, between DAS9 to DAS 12, the rapid growth of adventitious roots caused a temporary increase in NH_4_^+^ concentration in the roots. Despite this, as biomass, particularly shoot weight, increased rapidly, NH_4_^+^ in roots was quickly assimilated into various amino acids and enters the plant’s N metabolic process, resulting another decrease in NH_4_^+^ concentration in the roots. In comparison to LN, HN exhibited a significantly higher root fresh weight per plant at DAS 12 and DAS 15, with increases of 15.3% and 25.0%, respectively (Fig 2c, 2d). Additionally, the shoot-to-root weight ratios at DAS 9, DAS 12, and DAS 15 were 4.95%, 6.01%, and 8.87% higher in HN than in LN. Nevertheless, the rapid increase in biomass and NH_4_^+^ assimilation somewhat diluted the NH_4_^+^ concentrations in the roots, leading to lower root NH_4_^+^ concentrations in HN compared to LN during DAS 9−15 (Fig 2a).

**Fig. 2.**
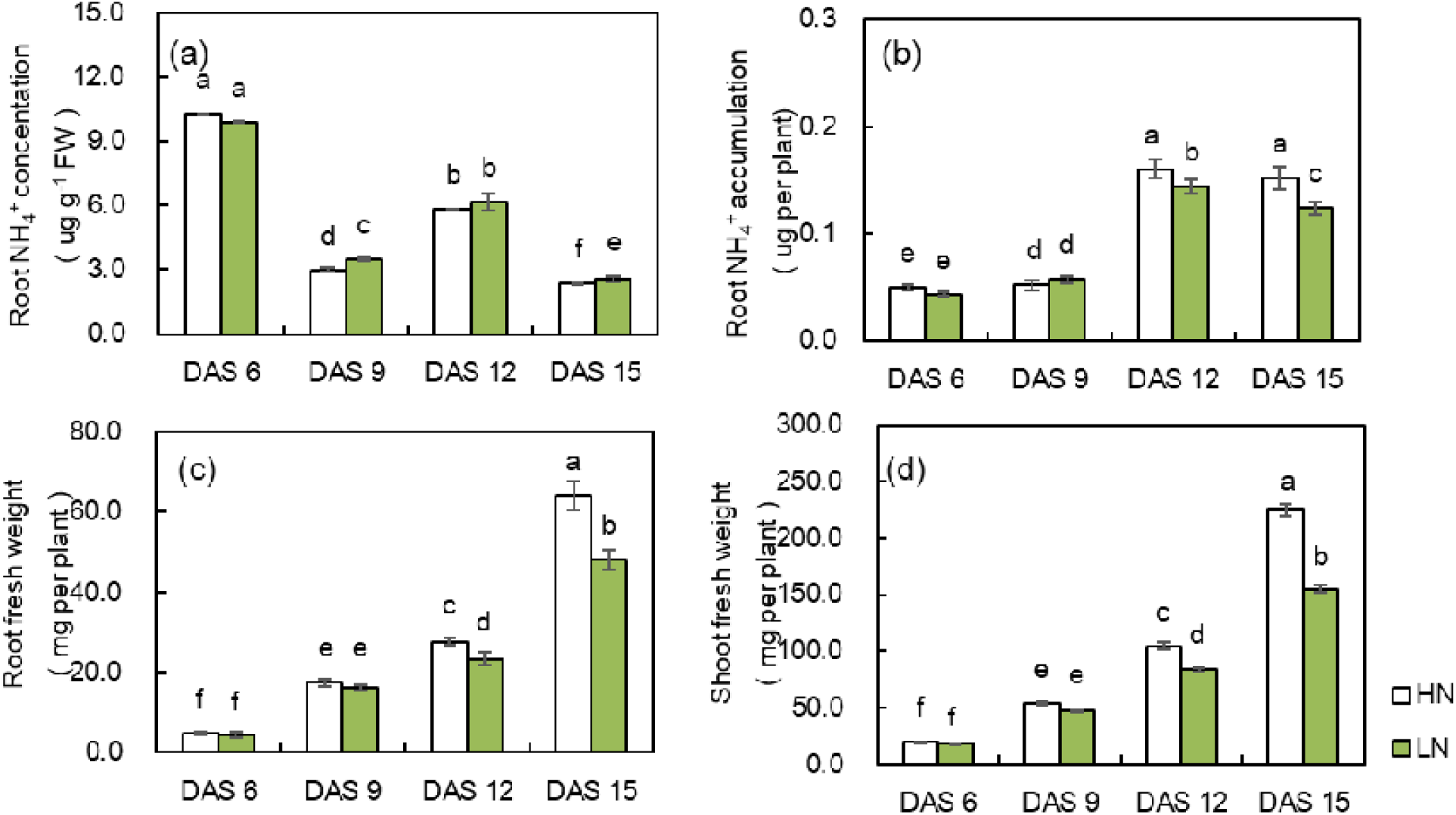
Root NH_4_^+^ concentration, accumulation and the fresh weights of both roots and shoots in rice seedlings under of HN and LN treatment.

### NH_4_^+^ flux at different root zones of the seminal root

NH_4_^+^ flux was investigated in the root cap, meristematic, elongation, maturation zones of the seminal root at DAS 6, 9, 12, and 15. As shown in Fig. 4, under both HN and LN treatments, the net NH_4_^+^ flux rates were negative in all root zones on DAS 6, indicating net NH_4_^+^ influx. On DAS 12, only the maturation zone and root hair region showed negative net NH_4_^+^ flux rates, while the root cap, meristematic, and elongation zones displayed positive rates, suggesting a net efflux of NH_4_^+^ from these three zones; By DAS15, NH_4_^+^ efflux was observed in all root zones, indicating that the entire seminal root had ceased its nutrient uptake function by this point.

**Fig. 3.**
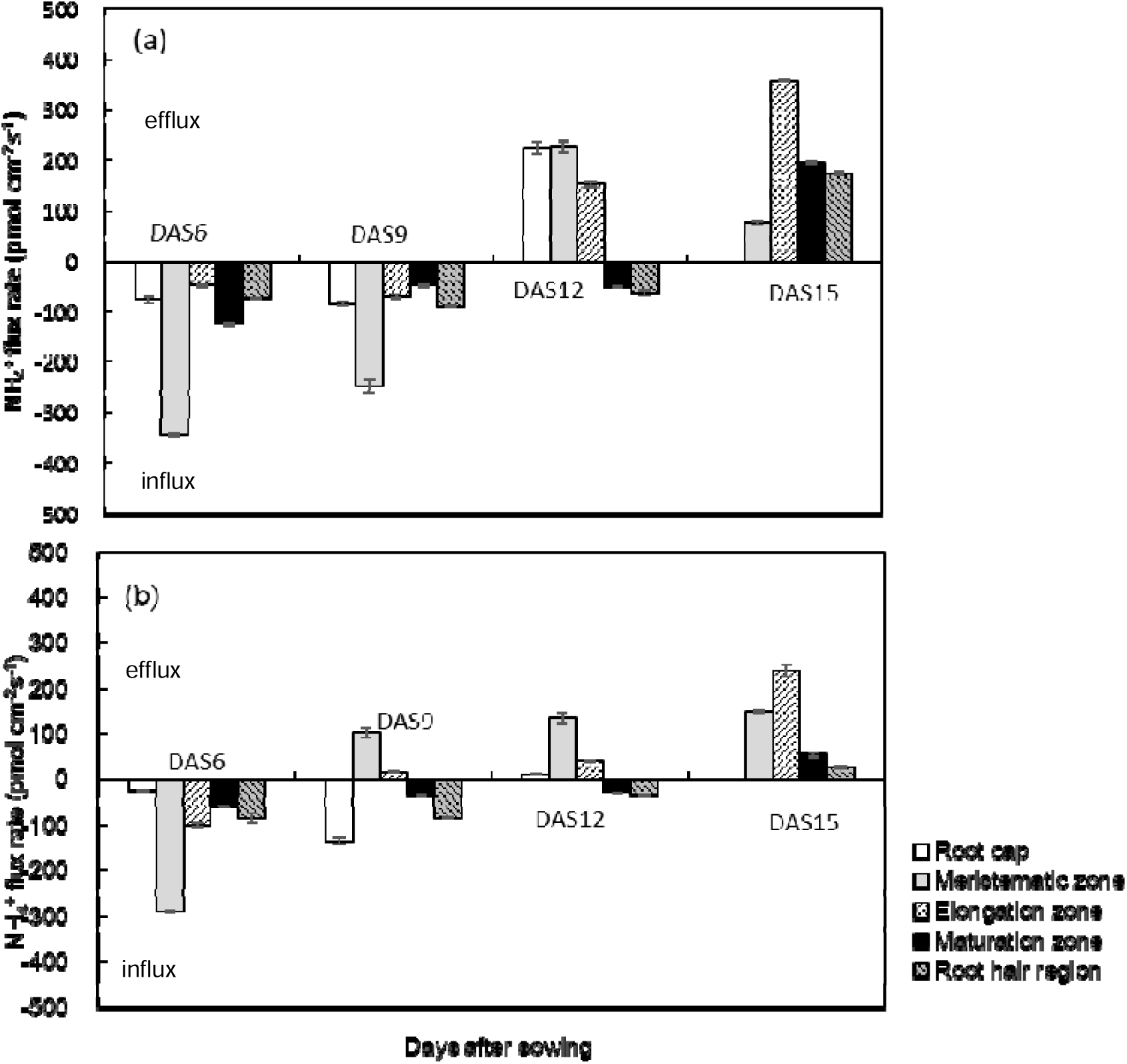
NH_4_^+^ flux in the root cap, meristematic, elongation, maturation zones of the seminal root at DAS 6, 9, 12, and 15. Positive values indicate net NH_4_^+^ efflux, negative values indicate net NH_4_^+^ influx.

**Fig. 4.**
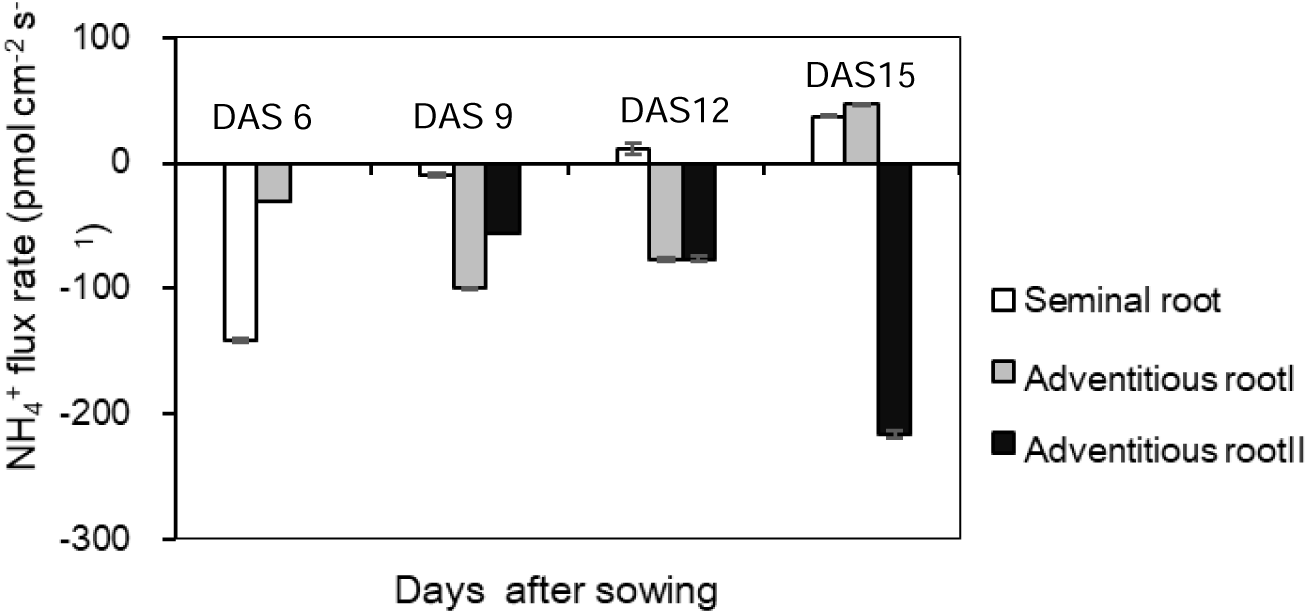
The NH_4_^+^ flux rates in the root hair region of the seminal and adventitious roots at DAS 6, DAS 9, DAS 12 and DAS 15

The meristematic zone was the primary site for nutrient uptake in new roots. During DAS 6 to DAS 9 in the HN substrate, the average net flux rates of NH_4_^+^ in the root cap, meristematic zone, elongation zone, and mature zone were -80 pmol cm^−2^ s^−1^, -285 pmol cm^−2^ s^−1^, -65 pmol cm^−2^ s^−1^, and -77 pmol cm^−2^ s^−1^, respectively, the net influx rate of NH_4_^+^ in the meristematic zone was significantly higher than in other root zones (Fig. 5a). In the LN substrate, the net flux rate of NH_4_^+^ in the meristematic zone at DAS 6 was -290 pmol cm^−2^ s^−1^, which was 10.8, 2.9, 5.0 times higher than the value of the root cap, elongation and mature zones, respectively (Fig.5b).

The maturation zone and root hairs were the main region for nutrient uptake in mature and aging roots. Unlike the root cap, meristematic, and elongation zones that experienced a net NH_4_^+^ efflux on DAS 9 or DAS 12, net NH_4_^+^ efflux was not detected in the maturation zone and root hairs until DAS 15 (Fig. 5), indicating both the maturation zone and root hair region had a nutrient uptake function for at least 12 days, significantly longer in time than root cap, meristematic, and elongation zones.

The NH_4_^+^ flux rate of roots was influenced by the external NH_4_^+^ concentration, with the meristematic zone being the most sensitive to changes in the external NH_4_^+^ concentration. As shown in Fig. 5, net NH_4_^+^ influx at DAS6 was observed in all root zones of both HN and LN treatments, and the value was higher in HN than in LN at the root cap, meristematic tissue, and maturation zones, whereas lower in HN than in LN at the elongation zone and root hair region; external NH_4_^+^ concentration did not affect the direction of NH_4_^+^ flux in the root cap and maturation zones, which changed from a net influx to a net efflux in both HN and LN treatments at DAS12 and DAS15, respectively; however, difference in the direction of NH_4_^+^ flux was observed in the meristematic and elongation zones at DAS9, with influx in the HN while efflux in the LN, and the meristematic and elongation zones in HN did not exhibit net NH_4_^+^ efflux until DAS 12, that is, increasing the external NH_4_^+^ concentration could prolong the time of net NH_4_^+^ influx in the meristematic and elongation zones by 3 days. At DAS 9, the ammonium ion flux rates in the meristematic zone under HN and LN were -247.6 pmol cm^−2^ s^−1^ (net influx) and 102.5 pmol cm^−2^ s^−1^ (net efflux), respectively, with a difference up to 350 pmol cm^−2^ s^−1^, which was much higher than the difference of 83.7 pmol cm^−2^ s^−1^ between HN and LN in the elongation zone.

### The process of new roots replacing old roots for ammonium absorption

Fig. 6 was the NH_4_^+^ flux rates in the root hair region of the seminal and adventitious roots at DAS 6, DAS 9, DAS 12 and DAS 15, it showed that the uptake of NH_4_^+^ by roots during the growth of rice seedlings was gradually transferred from the seminal root to adventitious root I, then to adventitious root II. The NH_4_^+^ flux rates in the root hair region of the seminal root gradually decreased, it was as high as -141.6 pmol cm^−2^ s^−1^ at DAS 6, and decreased to -9.78 pmol cm^−2^ s^−1^ at DAS 9; while that of adventitious root I gradually increased from -30.31 pmol cm^−2^ s^−1^ at DAS 6 to -100.08 pmol cm^−2^ s^−1^ at DAS 9, and that of adventitious root II gradually increased from -56.73 pmol cm^−2^ s^−1^ at DAS 9 to -216.08 pmol cm^−2^ s^−1^ at DAS 15. Changes in the rate of NH_4_^+^ flux in the root hair zone implied the process of new adventitious root gradually replacing the aging roots in the uptake of NH_4_^+^.

## Discussion

As the major organ for plant nutrient uptake and growth support, root morphological optimization and functional enhancement are of great importance for breakthroughs in yield and fertilizer use efficiency. The morphology characteristics of rice root include root number, root length, root radius, root surface area, and root density and so on. Typically, rice plants with deep and wide root distribution, as well as well-developed root hairs, exhibit a strong nutrient uptake capacity due to their larger contact surface area with the soil, while plants with shallow and small root have a weaker nutrient absorption capacity. However, it is very difficult to directly observe and analyze the root system growing in the soil. To overcome this, various methods have been employed by researchers, root morphology is monitored using methods such as excavation, localization, nail board, earth drill, wall surface and glass wall methods, and container techniques, etc., and root nutrient uptake capacity is measured by physiological and biochemical indicators, such as active absorption surface area, root activity, root respiration intensity and relative enzyme activity, etc. However, these methods are not only time-consuming and labor-intensive, but are also affected by the representativeness of the root samples. Non-invasive micro-test technology (NMT) can obtain the transmembrane flux rates of Ca_2_^+^, H^+^, K^+^, Na^+^, NH_4_^+^, NO_3_^−^ and other ions by microsensors without damaging the biological samples, which provides a method to finely analyze the NH_4_^+^ uptake capacity of the rice root system at a microscopic level (Xu *et al*., 2016).

Studies have shown that the absorption of NH_4_^+^ by rice roots exhibits dual kinetic characteristics. When the concentration of NH_4_^+^ is low (at the uM level), the high affinity transport system plays a dominant role and exhibits saturation kinetic characteristics; when the external NH_4_^+^ concentration is high (at the mM level), the low-affinity transport system playing a dominant role and does not show the characteristics of saturation kinetics (Wand *et al*., 1993; von Wiren *et al*., 2000; Britto *et al*., 2001), that is, under high NH_4_^+^ concentration, there is no down-regulation for NH_4_^+^ uptake in plants, and the higher the eternal NH_4_^+^ concentration, the more NH_4_^+^ influx, which is very likely to lead to excessive uptake of NH_4_^+^, and a large amount of NH_4_^+^ accumulates in the roots, some of which was assimilated into various amino acids and enters the N metabolic process in the plant (Kiyomiya *et al*., 2001; Xu *et al*., 2016), while excessive NH_4_^+^ was actively excreted out the root after NH_4_^+^ assimilation in roots reaching saturation and transmembrane fluxes of NH_4_^+^ was observed in roots (Chen *et al*., 2017). Britto *et al*. (2001) found that in the high NH_4_^+^ environment, there is a significant efflux of NH_4_^+^ in barley roots, and the amount of the efflux can even account for 80% of the initial NH_4_^+^ influx. Chen *et al*. (2013) investigated the NH_4_^+^ flux of rice root and found that the root meristematic zone was the main transmembrane influx site of NH_4_^+^, while the transmembrane efflux of NH_4_^+^ was mainly in the elongation zone of the root, and by increasing the external NH_4_^+^ concentration, the NH_4_^+^ efflux in the root elongation zone of the N-inefficient rice varieties was significantly increased. Our study further studied the transmembrane fluxes of NH_4_^+^ in different rice root zones, and found that the main regions of NH_4_^+^ influx and efflux were varied with root aging. In new roots, all root zones had transmembrane NH_4_^+^ influx with the meristematic zone had the greatest value, this may be due to active cellular metabolism in the meristematic zone in the new root, which favors NH_4_^+^ assimilation, while the Kjeldahl band is not yet fully formed and does not impede the transport of assimilated amino acids(Wang *et al*., 2014); In aging roots (9−12 days after root emergence), there was a net influx of NH_4_^+^ only at the mature and root hair zone, which was consistent with the study of Zhang *et al*. (2018), who found that the cell intercellular space in the root hair could act as a channels for nutrient transport and exhibited significant ATPase activity, and due to the dense distribution and large quantity of root hairs, the maturation zone where root hairs grow became the most vigorous region for water and nutrient absorption in robust roots (Gilroy and Jones, 2000; Leitner *et al*., 2010); while a net efflux of NH_4_^+^ was observed from the root cap, meristematic zone, and elongation zones probably due to the decrease in NH_4_^+^ assimilatory enzyme activity during cell senescence. However, in aging root 15 days after root emergence, the net transmembrane NH_4_^+^ influx in root hair region was also disappeared, and NH_4_^+^ efflux was observed across all root zones. In the supplementary experiment, we excised an aging seminal root and an aging adventitious root from the seedlings and measured NH_4_^+^ flux of the hair region of the excised roots. Both roots exhibited the same trend in NH_4_^+^ flux changes: transmembrane NH_4_^+^ efflux was observed even in the excised roots, but the efflux rate gradually decreased, ultimately dropping to zero after 60 hours of excision (supplementary Fig.3). We speculated that this might be due to the active excreted out of excessive NH_4_^+^ accumulated in the excised roots, which continued until the root cells gradually died and could no longer supply the energy required for NH_4_^+^ efflux.

In this study, we further investigated the differences in NH_4_^+^ flux in the roots of seedlings grow in HN and LN seedling-raising mediums, and found that the meristematic zone was the most sensitive to external NH_4_^+^ concentration. Rice seedlings grown in HN showed a significantly higher net NH_4_^+^ influx rate in the meristematic zone on DAS 6 compared to those grown in LN. Additionally, the meristematic zone in HN did not show a net NH_4_^+^ influx until DAS 12, which was a 3 days later than that at DAS 9 in LN.

During the growth of rice seedlings, new roots continuously emerge while older roots gradually die. The seminal root plays a crucial role only shortly after seed germination, its nutrient uptake function diminishes and disappeared after adventitious roots appear (Meng *et al*., 2019; Sun *et al*., 2022). Our microscopic observations revealed that as root age, the root cap gradually detached from the root tip, and the root tip changed from sharp to round, and the distance between the root hair and the root tip shortened. Further measurements of NH_4_^+^ flow using NMT in both new and aging roots revealed that the NH_4_^+^ uptake capacity of rice seminal and adventitious roots disappeared 12−15 days after root emergence. Meanwhile, new roots gradually took over the main nutrient uptake role in robust seedlings.

## Conclusions

A root can sustain NH_4_^+^ uptake capacity for approximately 12 days. As the root ages, the root cap gradually degraded from the tip, causing the distance between the maturation zone and the root tip to shorten. The meristematic zone was the primary region for NH_4_^+^ absorption in new roots and is highly sensitive to changes in the external N concentration, while the root hair region is the main site for NH_4_^+^ absorption in mature and older roots (9−12 days after root emergence). Throughout rice growth, new adventitious roots gradually replace seminal root and older adventitious roots as the primary force of nutrient uptake.

## Acknowledgements

This research was supported by grants from the National Natural Science Foundation of China (No. 32272210), and the National Rice Industry Technology System (CARS-01-31).

## Author contributions

D.W. designed and supervised the research. X.Z. conducted the experiments. S.C., G.C., C.X., and Y.Y. performed data analysis. X.Z. and Y.X. wrote the manuscript. All authors have read and approved the manuscript.

## Data availability

The data generated in the current study are available from the corresponding author on reasonable request.

